# OmicsResonance: An LLM-assisted cloud ecosystem for interactive and reproducible single-cell transcriptomic analysis

**DOI:** 10.64898/2026.09.20.753054

**Authors:** Li Chenhui, Qin Zhizhen, Lu Yi, Wang Qinfen, Chen Yan-kai, Li Yi, Zeng Lin, Chen Mingjie

## Abstract

While single-cell RNA sequencing (scRNA-seq) is indispensable, existing analytical tools impose high computational barriers, requiring complex environment configurations and programming proficiency. To democratize single-cell analysis, we developed OmicsResonance, a code-free, web-based platform that eliminates local installations and empowers researchers to execute end-to-end analyses directly within a browser. The platform integrates standard processing pipelines and Large Language Model (LLM)-assisted cell annotation with a suite of advanced modules, including pseudo-time inference, CNV profiling, and virtual knockout simulations. Built on a scalable architecture, OmicsResonance champions “justified analysis” by avoiding black-box defaults. Its interactive interface allows users to dynamically adjust parameters and instantly visualize biological impacts, ensuring transparent and biologically sound analytical decisions. Validation across three public datasets successfully reproduced established findings and facilitated further interpretation of the data. Ultimately, OmicsResonance bridges the gap between complex computational pipelines and bench research, providing an accessible yet powerful environment for scRNA-seq data exploration. The platform is available at https://cloud.rnastar.com/.

**Abstract Graph:** 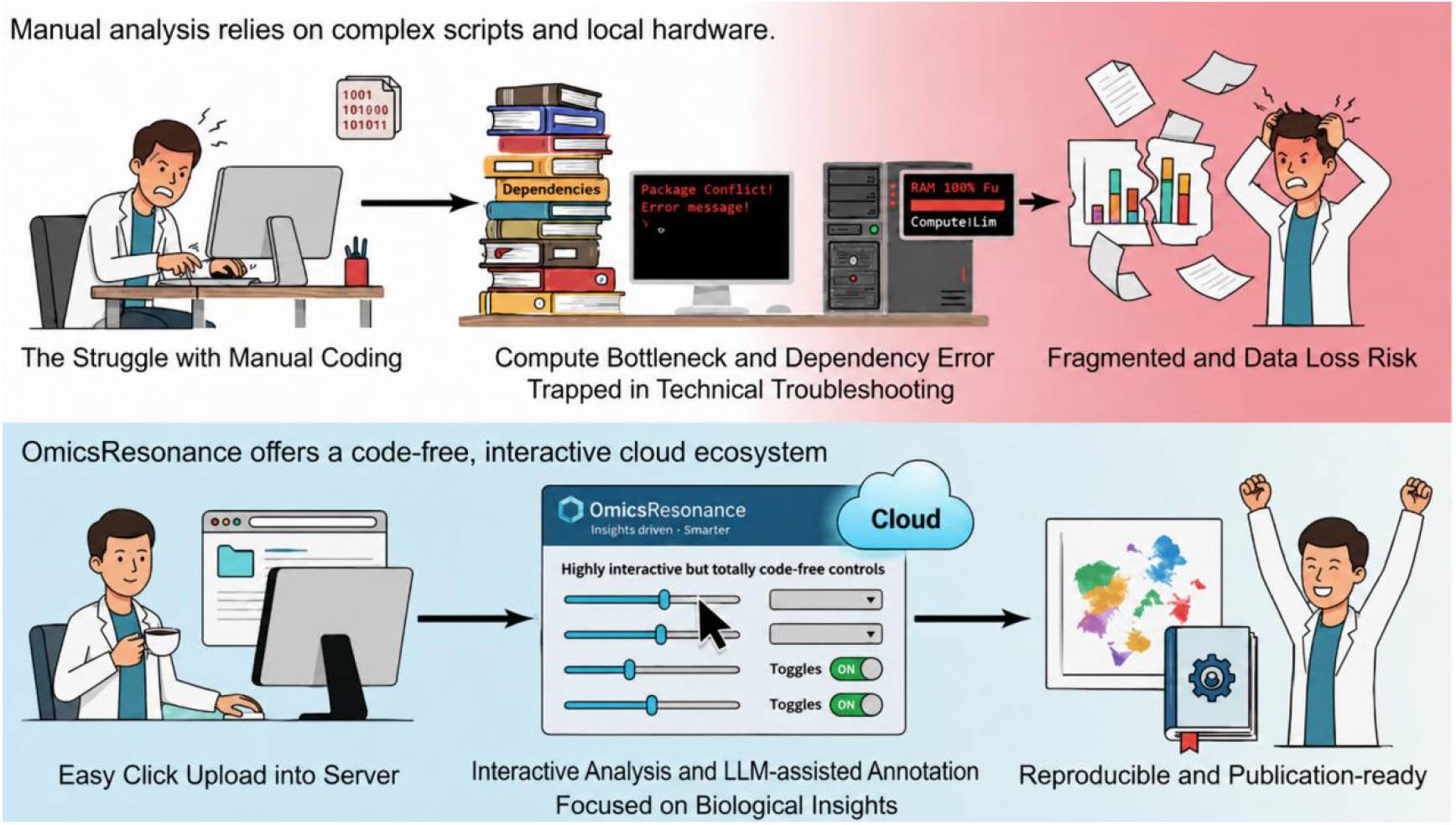

## Introduction

The advent of single-cell RNA sequencing (scRNA-seq) has fundamentally transformed life science research. Researchers can now dissect tissue heterogeneity[1, 2], delineate cellular developmental trajectories[3], and characterize complex disease micro-environments[4] at single-cell resolution. This paradigm shift has significantly advanced diverse fields, including developmental biology, tumor immunology, and regenerative medicine. Concurrently, the increasing accessibility of sequencing platforms has generated massive amounts of transcriptomic data, stimulating a rapid proliferation of specialized bioinformatics tools. Despite these methodological advancements, the transition from raw data generation to biological insight remains a substantial bottleneck. The continuous emergence of analytical software has inadvertently introduced a new set of logistical and technical challenges. Specifically, researchers attempting to navigate this ecosystem encounter four primary obstacles.

First, the proliferation of over a thousand available scRNA-seq tools[5] complicates workflow standardization. These disjointed analytical workflows force researchers to spend disproportionate time benchmarking algorithms rather than focusing on downstream biological interpretation. Second, traditional workflows impose severe technical and infrastructural barriers. Most tools require rigorous programming proficiency and intricate dependency management, frequently causing version conflicts that compromise reproducibility. Furthermore, the exponential growth of scRNA-seq datasets routinely exhausts local memory limits, forcing reliance on specialized high-performance computing facilities. Third, a significant disconnect persists between computational tool design and biological end-users. While recent large language models (LLMs)[6] and autonomous agents[7, 8] partially lower coding barriers, they introduce new, opaque challenges. AI can rapidly generate preliminary outputs, but these often lack transparency and adequate biological context. Consequently, researchers must still invest substantial effort to manually verify whether these AI-generated insights are biologically sound. Finally, the predominant reliance on command-line interfaces creates a critical accessibility barrier. The lack of intuitive interactive interfaces limits immediate visual feedback during parameter adjustment, restricting biological researchers from dynamically exploring and interrogating their data.

In response to these obstacles, developing an integrated, online one-stop analysis platform has emerged as an ideal solution. Several online platforms have been released, such as ICARUS[9], BestopCloud[10], CytoAnalyst[11], SingleCAnalyzer[12], and scExplorer[13]. However, these platforms generally suffer from several limitations that restrict their widespread use. For example, scExplorer[13], despite its user-friendly interface and guidance on analysis parameters and steps, exhibits a confusing analytical workflow: it allows users to enter the annotation interface before marker genes have been calculated, and the annotation interface does not display marker genes, making it difficult for researchers to effectively utilize the platform. CytoAnalyst[11] lacks user guidance, imposes restrictions on the sources of analyzable data (some platform data cannot be used), and provides non-functional demo data, preventing researchers from testing platform functionality through examples. Moreover, existing platforms share common issues, such as failure to effectively remove doublets during quality control and a lack of correction methods for ambient RNA contamination. Finally, beyond these technical and functional deficits, a broader systemic problem is the lack of long-term sustainability; many of these web platforms become entirely inaccessible or cease to be maintained shortly after their corresponding papers are published[14, 15].

To address these limitations, we developed OmicsResonance, a web-based platform engineered within the R environment[16] to harness its robust statistical capabilities and extensive bioinformatics ecosystem. The analytical framework is anchored by Seurat v5.3.0[17] and systematically integrates a suite of established analytical methods together with an LLM-assisted component for supporting cell annotation. To eliminate programming barriers, OmicsResonance features an intuitive graphical user interface (GUI) enriched with contextual documentation. Interactive tool tips and inline explanations for all adjustable parameters allow users to evaluate the biological impact of each analytical choice in real time. By abstracting complex computational pipelines into this transparent, parameter-driven framework, researchers can autonomously execute end-to-end scRNA-seq analyses, ensuring a rigorous transition from raw data processing to biological interpretation.

OmicsResonance transitions from standard baseline processing to a comprehensive suite of advanced downstream modules, facilitating complex explorations such as pseudo-time inference, cell communication, and virtual knockout simulations. To resolve technical noise and batch effects, the platform deliberately avoids rigid, “black-box” automated pipelines in favor of a transparent, user-driven iterative workflow [18, 19]. Empowered by contextual tool tips and real-time visual feedback, researchers can dynamically tune parameters and confidently optimize data integration strategies for their specific datasets. This principle of “justified analysis” extends to cell annotation, where database-driven[20–24] and LLM-assisted predictions serve as an intelligent baseline that biologists can rigorously refine through transparent marker gene visualization. Ultimately, OmicsResonance bridges the gap between high accessibility and analytical depth, empowering bench scientists to independently translate high-dimensional data into robust biological insights.

To evaluate the platform’s practical utility, we re-analyzed three independent public datasets across varying analytical complexities. First, a standard PBMC 3k dataset[25] validated the baseline workflow, demonstrating efficient quality control, dimensionality reduction, and accurate major cell type identification. Second, an intervertebral disc degeneration (IVDD) dataset[26, 27] highlighted our subpopulation refinement capabilities by successfully sub-clustering isolated nucleus pulposus cells into specific functional subgroups. Third, a complex mouse brain stroke (MCAO) dataset[28] requiring rigorous batch-effect correction showcased the seamless integration of advanced modules. By applying trajectory inference, stemness evaluation, and cell communication modeling to isolated endothelial cells, we confirmed the platform’s capacity for in-depth biological exploration. Collectively, these analyses not only reproduced established findings but also facilitated deeper interpretation of the data. To facilitate user adoption, comprehensive step-by-step video tutorials for all three workflows are provided, reaffirming OmicsResonance as a robust and highly accessible tool for single-cell research.

## Material and Methods

### Data Import

Given the bandwidth and storage constraints of web-based applications, OmicsResonance exclusively processes post-alignment expression matrices; the upload of raw sequencing data (e.g., FASTQ files) is not supported due to prohibitive file sizes. The system accommodates diverse matrix-level inputs from major platforms (e.g., 10X Genomics, MGI), encompassing standard 10X output directories (barcodes.tsv, features.tsv, and matrix.mtx), single-file formats (.h5, .h5ad), and standalone count matrices. To streamline data transfer, users can upload standard compressed archives (e.g., ZIP, RAR, TAR). Upon upload, the platform automatically identifies the file format, extracts the contents, and validates the data structure, eliminating manual configuration. Detailed formatting guidelines and downloadable example datasets are readily accessible on the platform’s data upload interface.

### Quality Control

OmicsResonance features an interactive quality control (QC) module that enables researchers to iteratively adjust filtering thresholds based on statistical distributions and biological context, effectively clearing technical noise while preserving rare, meaningful cell populations. Notably, this real-time, visual “adjust-and-re-run” design principle is universally implemented across all platform modules and will not be reiterated in subsequent sections.

The QC workflow leverages standard Seurat-derived metrics[29]. Users can define cutoffs for detected genes (nGene) and UMI counts to exclude empty droplets and potential multiplets, and utilize the log10(GenesPerUMI) metric to evaluate sequencing complexity. Furthermore, the module incorporates mitochondrial, ribosomal, and hemoglobin ratios to flag apoptotic cells or unwanted blood contamination, alongside advanced options for ambient RNA removal, manual doublet filtering, and minimum cell-expression thresholds. To establish a baseline, the platform provides empirical default parameters: nGene ranging from 250 to 8,000; UMI counts from 500 to 15,000; a minimum log10(GenesPerUMI) of 0.8; and a maximum mitochondrial ratio of 20%. By default, ribosomal and hemoglobin ratios are unbounded (capped at 100%), ambient RNA removal is disabled, and genes must be expressed in at least 10 cells. Guided by real-time visualizations, users can dynamically fine-tune these parameters to accommodate sample-specific characteristics.

### Sample Integration and Dimension Reduction

Following quality control, OmicsResonance provides flexible modules for data regression, integration, and dimensionality reduction. To ensure clustering is driven by genuine biological identity rather than technical bias, users can selectively regress out confounding variables, including total UMI counts, mitochondrial ratios, and cell cycle effects. By default, only the mitochondrial ratio is regressed out to mitigate metabolic stress artifacts, while UMI and cell cycle regressions are disabled to prevent over-correction of genuine biological variance. For multi-sample datasets, the platform addresses batch effects by offering multiple robust integration algorithms[18, 19, 30], enabling researchers to seamlessly align data across different donors or experimental conditions. Finally, users retain precise control over feature selection and dimensionality reduction by defining the number of highly variable genes (HVGs) and principal components (PCs), which are preset to empirical defaults of 3,000 genes and 50 components, respectively. This customizable parameterization ensures the final dimensional space optimally captures both major cellular lineages and rare subpopulations.

### Cell Clustering and Marker Gene Identification

OmicsResonance provides precise control over cell clustering and marker gene extraction. Users can specify the number of principal components (PCs) and adjust the clustering resolution to dynamically dictate the granularity of cell state detection, ranging from broad lineages to rare transient states. For subsequent marker gene computation, researchers can define stringency thresholds, specifically, the minimum log fold-change and the minimum percentage of expressing cells (min.pct), to ensure robust identification of subpopulation-specific signatures. Furthermore, users can trigger an optional automated enrichment analysis to instantly map these computed markers to established biological pathways. The module is initialized with empirical defaults: 30 PCs, a clustering resolution of 1.0, a minimum log fold-change of 0.1, and a min.pct of 0.1. By iteratively fine-tuning these exact parameters, users can accurately transform mathematical clusters into verified biological entities.

### Cell Annotation

Cell type annotation remains a critical yet labor-intensive bottleneck in single-cell transcriptomics. To address this, OmicsResonance streamlines the process by integrating predictions from established algorithms, LLM-assisted interpretation, and authoritative reference databases into a cohesive, biologically justified workflow. First, the platform deploys an ensemble of five established automated annotation algorithms (SingleR[20], SCINA[21], CellID[22], ScType[24], and scCATCH[23]). Rather than presenting disjointed outputs, the system synthesizes these results within an interactive dashboard. This interface provides comprehensive diagnostic visualizations, robust population statistics (counts and percentages), and transparent marker gene expression mapping. Crucially, researchers can dynamically review cluster-specific markers alongside algorithmic assignments, empowering them to manually override or refine labels to ensure biological accuracy.

Complementing the conventional annotation methods, OmicsResonance uses DeepSeek-V4-pro[31] as an auxiliary evidence-integration component rather than as a standalone annotation algorithm. For each annotation task, the backend constructs two structured inputs: a cluster-by-method table containing cluster-level predictions from the available conventional annotation methods, and a cluster-specific marker table containing the retained marker genes and their associated statistics. Marker records passing the platform’s predefined filtering criteria are ranked within each cluster, and up to 200 records per cluster, together with their original statistical fields and column headers, are included. Information from all clusters is combined into a single structured JSON request.

The request contains a system message instructing the model to act as a bioinformatics specialist and return an English JSON response, and a user message describing the two input tables and requesting a candidate cell-type label and supporting reason for each cluster. DeepSeek-V4-pro[31] was used with a temperature of 0.7 and JSON-object response mode; other generation parameters followed the API defaults. The expected output is a cluster-keyed JSON object containing “celltype” and “reason” fields. The returned suggestions are displayed in the annotation interface without automatically replacing the final labels. Users compare these suggestions with the conventional assignments and marker-gene evidence, consult CellMarker 2.0[32] separately, and manually confirm or revise the final annotation. CellMarker 2.0[32] is used for manual validation and is not directly included in the LLM prompt.

### Evaluation of LLM-assisted annotation

We compared the LLM-assisted annotation output with SingleR[20], SCINA[21], CellID[22], ScType[24], and scCATCH[23] across PBMC 3K[25], GSE174574[28], and GSE244889[26, 27]. Manually curated cell-type annotations from the corresponding source studies or the Seurat PBMC 3K tutorial, supported by their reported marker genes, were used as reference labels. Accuracy was calculated from cluster-level agreement, whereas precision, recall, and F1 score were macro-averaged across reference cell-type categories. For PBMC 3K, the marker-gene list required by SCINA was derived from the canonical markers in the Seurat PBMC 3K tutorial. SCINA was not evaluated for GSE174574[28] because no equivalent predefined marker list covering its expected cell types was available without introducing additional prior cell-type judgment. Each dataset was evaluated using one LLM-assisted output; repeated independent generations were not performed.

### Optional Personalized Analyses

To facilitate in-depth mechanistic exploration beyond baseline profiling, OmicsResonance integrates a comprehensive suite of advanced functional modules. The platform directly empowers users to execute rigorous differential expression analysis (Wilcoxon test) and compute pathway or gene set enrichment scores (AUCell[33], AddModuleScore[34], UCell[35]). For resolving dynamic cellular processes, researchers can map developmental trajectories (Monocle3[36]), evaluate differentiation states (CytoTRACE2_1.1.0[37]), and model intercellular communication networks (CellChat[38]). Additionally, the system supports complex genomic and perturbational interrogations, including large-scale chromosomal profiling (InferCNV[39]) and virtual genetic knockout simulations (ScTenifoldKnk[40]). Beyond this current analytical repertoire, OmicsResonance is engineered with a scalable, highly extensible architecture. This modular framework facilitates the rapid integration of emerging computational algorithms, ensuring the platform continuously evolves to meet the advancing demands of the single-cell research community.

### Platform Architecture

OmicsResonance architecture is built upon a robust LNMP stack. For the user interface, the Vue 3.0 framework delivers highly responsive and interactive web components. To connect this front-end to the computational engines, a ThinkPHP 6.0 framework provides a secure RESTful API. Crucially, the platform uses Server-Sent Events (SSE) to push real-time task progress directly to the client. This ensures users can monitor long-running analyses without experiencing browser timeouts. At the computational core, the server orchestrates decoupled R and Python scripts via a custom job queue. Standard workflows rely on the Seurat ecosystem(v5.3.0)[17], while advanced modules operate independently. This modular design makes the system highly extensible, allowing new bioinformatics algorithms to be integrated easily in the future. The platform is deployed on a dual-socket HPC server equipped with two AMD EPYC 9654 96-core processors, providing 384 logical CPUs in total, together with 792 GB RAM and 20 TB storage. This infrastructure supports concurrent analyses, although practical performance depends on dataset size, analytical workload, and available backend resources.

### Data security, privacy, and retention

Communication between users’ web browsers and the OmicsResonance server is encrypted in transit using HTTPS/TLS. User accounts, uploaded datasets, and analysis projects are logically isolated, and authentication and access-control mechanisms restrict access to corresponding projects and analytical resources. OmicsResonance follows a user-controlled retention model: uploaded datasets and generated results remain available until deleted by the corresponding user, and no fixed-duration automatic deletion policy is currently applied. Users are responsible for ensuring appropriate authorization, ethical approval, and de-identification before uploading human clinical data. The current centralized platform does not claim universal GDPR or HIPAA compliance, and the applicability of these frameworks must be evaluated for each deployment and use scenario.

### Reproducibility and version management

Analyses in OmicsResonance are executed within a centrally managed server-side software environment. Key software and model versions are controlled centrally, including the Seurat v5.3.0[17] analytical framework and DeepSeek-V4-pro[31] used for LLM-assisted annotation. User-defined parameters and execution logs are archived for individual projects. For LLM-assisted annotation, the selected model and generated output are recorded, and a previously stored output may be reused when a request matches the current cache key. Such cache reuse supports workflow-level consistency and traceability but does not demonstrate the intrinsic reproducibility of independently generated LLM responses. The current platform does not provide a user-facing containerized distribution, and updates to underlying software packages or models may influence analytical outputs.

### The methodology of reanalysis

To validate the platform’s diverse analytical capabilities, we executed three distinct workflows of increasing complexity. First, a standard PBMC 3k[25] dataset demonstrated the baseline pipeline. Following initial QC (retaining cells with 250-8,000 genes, 500-15,000 UMIs, < 20% mitochondrial ratio, and >0.8 novelty score), we processed 2,000 highly variable features (HVGs) using 10 principal components (PCs). Clustering (resolution = 0.5) and subsequent annotation via the CIBERSORT LM22[41] gene set successfully resolved major cell lineages. Second, an intervertebral disc degeneration (IVDD) dataset (GSE244889)[26, 27] highlighted our subpopulation refinement capabilities. After filtering, data scaling with UMI/mitochondrial regression, and global annotation via SingleR[20], we isolated the nucleus pulposus cell (NPC) population. Sub-clustering these NPCs (10 PCs, resolution = 0.8) successfully mapped distinct functional subsets, confirmed by canonical marker genes including CP, MSMO1, FBLN1, UBE2C, DKK1, and CHI3L2. Third, a multi-replicate mouse brain MCAO dataset (GSE174574)[28] showcased complex batch integration and advanced modular analysis. Following rigorous QC (<15% mitochondrial and <0.1% hemoglobin ratios) and Harmony-based batch correction across 2,000 HVGs, global populations were clustered (30 PCs, resolution = 1.0) and annotated using an ensemble approach (SingleR[20], ScType[24], scCATCH[23], and CellID[22]). We then isolated endothelial cells for Harmony-integrated sub-clustering (25 PCs, resolution = 0.7), seamlessly deploying advanced modules to execute gene set scoring, CytoTRACE2_1.1.0[37] stemness evaluation, Monocle[36] trajectory inference, and CellChat communication modeling[38].

To ensure strict computational reproducibility and facilitate rapid user adoption, comprehensive step-by-step video tutorials corresponding to all three analytical workflows are readily accessible on the platform.

## Result

### Overview of OmicsResonance single cell analysis platform

OmicsResonance provides a structured workflow for single-cell transcriptomic analysis, encompassing the entire process from data input to final report generation (Figure 1). The pipeline initiates with data upload and format validation. Subsequently, the expression matrices undergo rigorous quality control, batch effect integration, dimensionality reduction, and unsupervised clustering to establish a robust analytical baseline. Following this stage, researchers assign biological identities to the computational clusters. Given the inherent complexity of cell annotation, OmicsResonance employs a multi-layered, evidence-based approach. The platform generates initial predictions using an ensemble of five distinct automated algorithms and transparently cross-references cluster-specific marker genes with the authoritative CellMarker 2.0 database[32]. Building on this, an integrated LLM synthesizes these algorithmic outputs and marker profiles to provide context-aware annotation suggestions, complete with explicit biological reasoning. By presenting this comprehensive suite of computational and literature-based evidence, the platform empowers researchers to confidently cross-validate information and finalize cellular identities.

**Figure 1.**
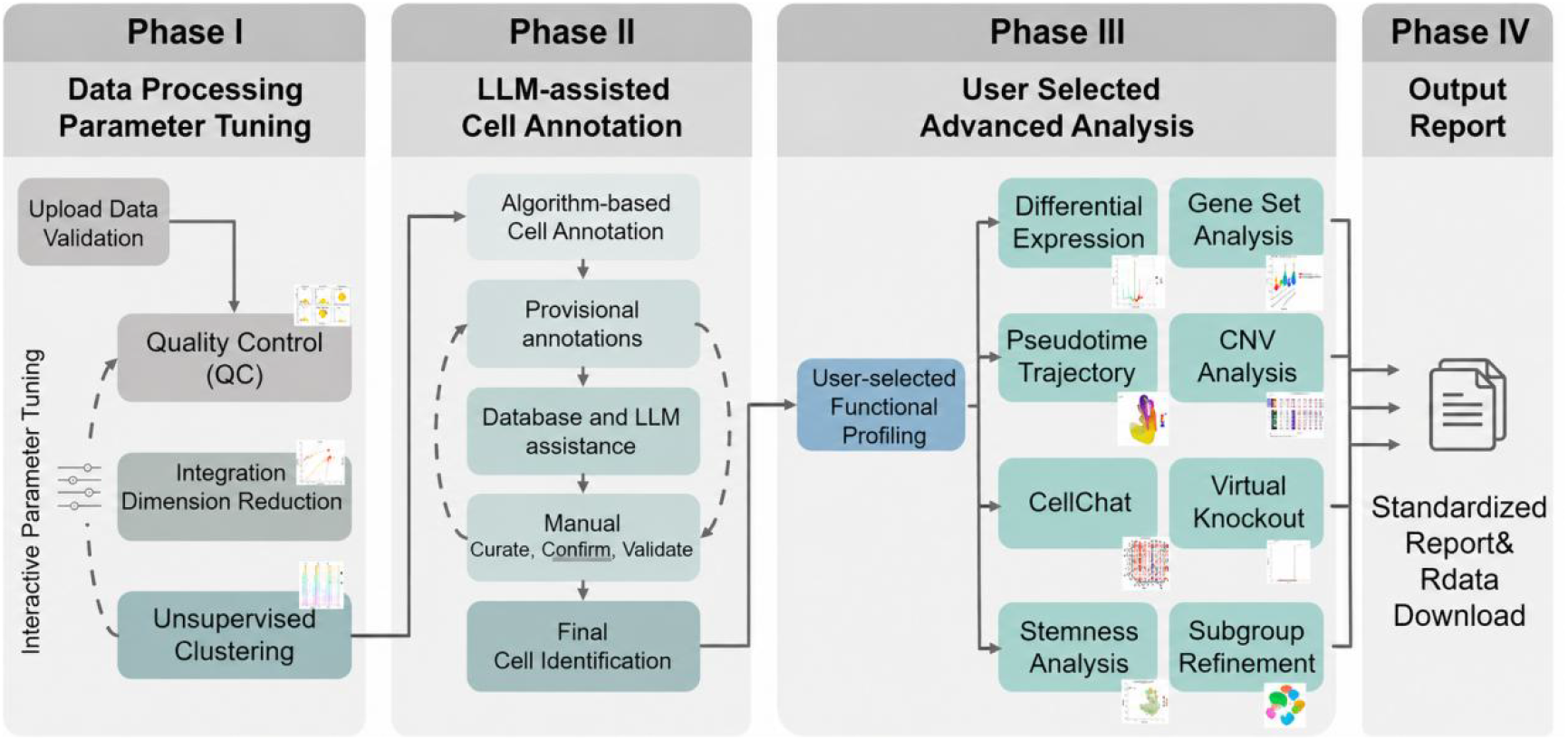
Overview of OmicsResonance single-cell analysis platform. The platform operates through four modular phases. Phase I handles data upload, quality control, and unsupervised clustering via interactive parameter tuning. Phase II executes a multi-method, LLM-assisted cell annotation workflow, integrating multi-algorithm predictions, database cross-referencing, and LLM reasoning to facilitate rigorous manual curation. Phase III empowers users to dynamically conduct subgroup refinement and select from a diverse suite of advanced functional profiling modules. Finally, Phase IV compiles all analytical outputs into standardized, publication-ready reports and downloadable .RData files for workflow traceability and reproducibility.

Once the cellular hierarchy is defined, the workflow transitions to hypothesis-driven functional exploration. In this phase, researchers can selectively deploy a comprehensive suite of advanced modules tailored to their specific questions. Crucially, this includes a subgroup refinement tool that allows users to dynamically tune clustering parameters to resolve and interrogate finer cellular states. Alongside this targeted sub-clustering, users can conduct differential expression comparisons and selectively deploy advanced modules, including gene set analysis, pseudo-time inference[36], copy number variation (CNV) profiling[39], cell-cell communication modeling[38], stemness evaluation[37], and virtual knockout simulations[40] based on their specific research questions.

Finally, the platform synthesizes all generated visualizations and analytical data into a standardized report for direct download. The following sections detail the specific implementation of these steps.

### Core Advantages and Comprehensive Capabilities of OmicsResonance

Compared to existing cloud-based platforms, OmicsResonance addresses prevalent analytical limitations through four practical advantages, focusing on analytical depth, quality control, and computational reproducibility (Table 1). First, broad accessibility and format compatibility. OmicsResonance overcomes standard input restrictions by automatically parsing diverse compressed archives and matrix formats (e.g., 10X, .h5ad, raw text). It offers comprehensive bilingual (English/Chinese) interfaces. This architecture eliminates local configuration hurdles, lowering the technical barrier for global researchers. Second, rigorous preprocessing and multi-method LLM-assisted annotation. Unlike many platforms that omit advanced noise reduction, OmicsResonance natively integrates ambient RNA cleanup and doublet removal. For cell identity assignment, it presents the outputs of five automated annotation methods together with cluster-specific marker evidence and an auxiliary LLM suggestion for user review. By synthesizing outputs from five automated algorithms[20–24], cross-referencing the CellMarker 2.0 database[32], and leveraging an integrated LLM to provide context-aware biological reasoning, this multi-layered approach supports transparent, user-reviewed annotation. Third, comprehensive analytics and cross-platform visualization. Beyond standard clustering, the platform incorporates specialized modules frequently absent in comparable tools, such as CNV profiling[39], trajectory inference[28, 36], and virtual knockout simulations (scTenifoldKnk)[40]. Furthermore, it supports .cloupe file export. This functionality allows researchers to conduct interactive, offline exploration within the Loupe Browser, independent of the original sequencing hardware. Finally, computational reproducibility and project administration. Single-cell workflows involve intricate, easily lost parameter settings. To support workflow reproducibility, OmicsResonance archives all user configurations, execution logs, and stoppable sessions. This enables complex analyses to be paused and resumed at will. Subsequently, a dedicated reporting module synthesizes these tracked workflows into publication-ready PDF reports, streamlining the transition from Cell Ranger-processed count matrices to manuscript preparation.

**Table 1.** Cross-comparison between OmicsResonance and other databases.

| Function/Platform | OmicsResonance | ICARUS v3 | BestopCloud | SingleCAnalyzer | scExplorer |
| --- | --- | --- | --- | --- | --- |
| Upload file format | rar, zip, tar.gz, gz | h5, h5ad, rds | gz | Fastq, h5 | h5ad, h5, rds, gz |
| Data format | 10x, h5, h5ad, txt | 10x, h5, h5ad, rds | gz, txt | 10x, h5 | h5ad, h5, rds, gz, 10x |
| Ambient RNA removal | Yes | No | No | No | No |
| Doublet removal | Yes | Yes | No | No | No |
| Celltype annotation | 5 methods, manual check and revision | 4 methods | 2 methods | SingleR only | No |
| LLM Assistance | Yes | No | No | No | No |
| Personalized analysis | Yes | No | No | No | No |
| Bilingual Support | EN/CN | EN | EN | EN | EN |
| Pausable sessions | Yes | No | No | No | No |
| .Rdata download | Yes | Yes | Yes | No | No |
| Project management | Yes | No | No | No | No |
| PDF report | Yes | No | No | No | Yes |

10x refers to the standard 10x files, including matrix.mtx, barcodes.tsv, and features.tsv (genes.tsv). "no" indicates that testers did not find relevant documentation under that platform, or did not find the relevant function during normal testing procedures. Personalized analysis includes differential expression analysis, stemness analysis, virtual knockout, pseudotime trajectory analysis, gene set analysis and cell communication analysis.

### Computational performance and scalability

We evaluated the computational performance of OmicsResonance using datasets containing approximately 5,000, 50,000, 100,000, and 500,000 cells. Each dataset size was benchmarked once under the same computational environment on a dual-socket server equipped with two AMD EPYC 9654 96-core processors, providing 384 logical CPUs in total with two hardware threads per physical core. Peak memory usage and total runtime were recorded (Table 2).

**Table 2.** Single-run benchmark of peak memory usage and total runtime of OmicsResonance for datasets with increasing numbers of cells under the same computational environment.

| Number of cells | Peak memory (GB) | Total runtime (min) |
| --- | --- | --- |
| 5,000 | 5.3 | 32 |
| 50,000 | 42.8 | 95 |
| 100,000 | 95.2 | 192 |
| 500,000 | 386 | 1,120 |

Both runtime and memory requirements increased substantially with dataset size. Although the 500,000-cell dataset was successfully processed in the tested environment, it required considerable high-memory computing resources. These results represent tested performance rather than a universal maximum supported dataset size.

### Validation of Fundamental Analytical Reliability Using PBMC Reference Data

To evaluate computational reliability, we replicated the 10X Genomics PBMC tutorial workflow. Applying identical quality control thresholds yielded diagnostic profiles consistent with the original dataset (Figure 2A, 2F; Supplementary Data S1). Subsequent dimensionality reduction generated UMAP topologies structurally comparable to the tutorial reference (Figure 2B, 2C); minor shifts in the UMAP layout reflect routine version updates in the underlying Seurat dependencies. For cell identity assignment, annotation via the LM22 reference set successfully recovered all major PBMC lineages. While minor differences in subpopulation labeling occurred (e.g., classifying “ActNK” as “NK cells resting,” Figure 2C) due to the use of alternative reference dictionaries, the core cellular hierarchy remained robust. Crucially, to mitigate the inherent subjectivity of automated labeling, we validated the expression patterns of canonical marker genes. The expression distributions of key myeloid markers, including *FCGR3A* and *MS4A7*, accurately reproduced the official tutorial (Figure 2D, 2E, 2G, 2H). This concordance not only validates the pipeline’s algorithmic accuracy but also reinforces the platform’s emphasis on transparent, visually guided manual verification of cellular identities.

**Figure 2.**
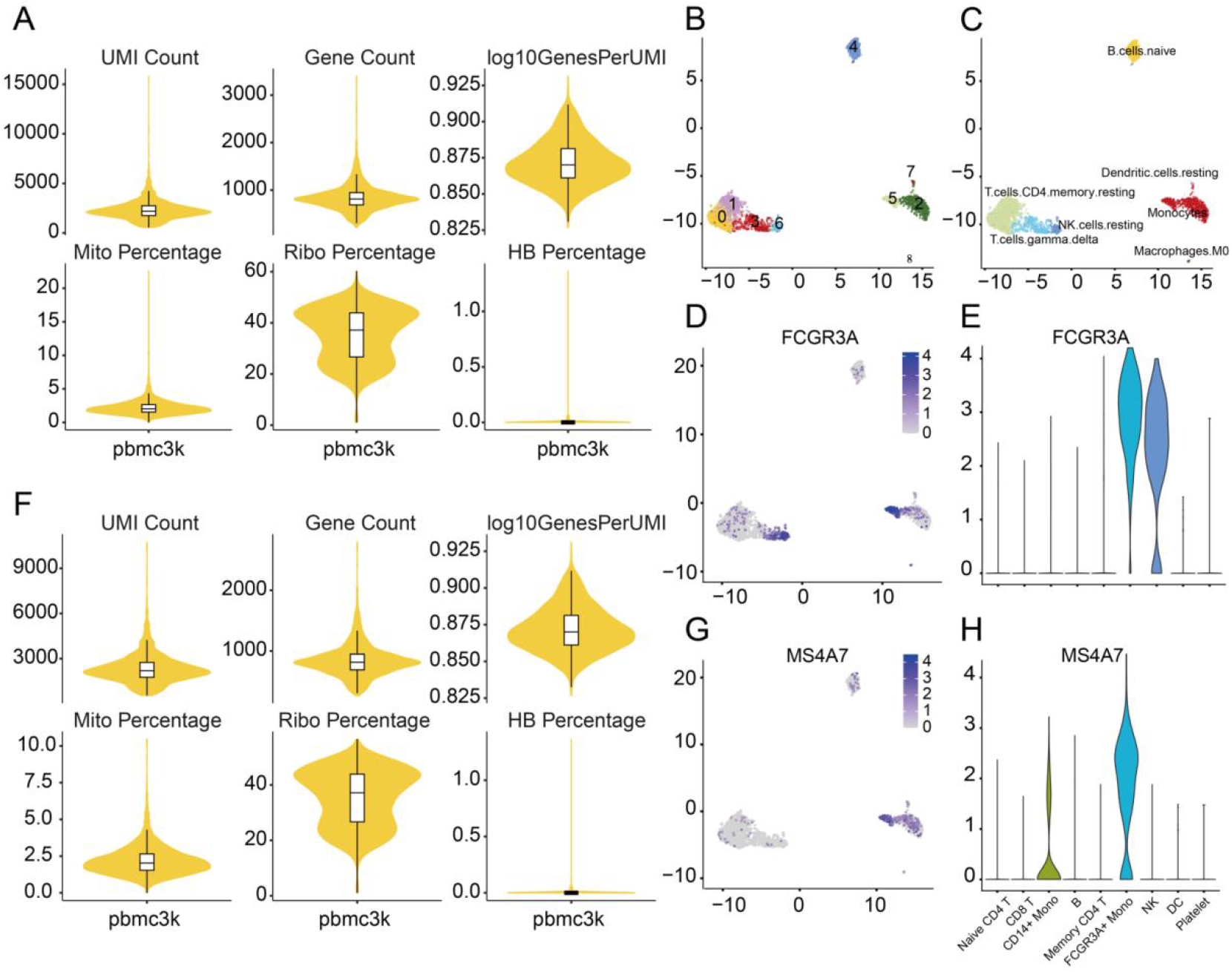
Validation of the baseline analytical workflow using the human PBMC 3k dataset. (A, F) Violin plots detailing the distribution of key quality control metrics before (A) and after (F) data filtering. (B, C) UMAP visualizations illustrating unsupervised cell clustering (B) and resulting global cell type annotations derived via CIBERSORT (C). (D-H) Expression profiles of representative subpopulation markers. UMAP feature plots display the expression patterns of *FCGR3A* (D) and *MS4A7* (G), with their relative expression levels across annotated cell types quantified in corresponding violin plots (E, H).

### High-Resolution Sub-Clustering and Heterogeneity Exploration

To evaluate the platform’s capacity for resolving complex tissue heterogeneity, we re-analyzed a published IVDD single-cell dataset comprising four mild (MDD) and three severe (SDD) nucleus pulposus samples [26, 27]. Following rigorous QC, 43,637 high-quality cells were clustered (resolution = 0.8) and annotated into eight major lineages, encompassing both immune (B/plasma, T cells, monocytes/macrophages) and non-immune (neutrophils, NPCs, endothelial cells, erythrocytes) compartments (Figure 3A). Notably, proportional analysis revealed a higher immune cell fraction in the MDD group compared to SDD (Figure 3B). While contrasting with the conventional expectation of heightened immune infiltration in severe degeneration, this accurately captures the inherent sample-level biological variance present in the original data. Furthermore, canonical marker expression profiles strictly aligned with established literature (Figure 3C), with the predominant NPC population robustly expressing *SOX9*, *ACAN*, and *PLOD2* (Figure 3D-F).

**Figure 3.**
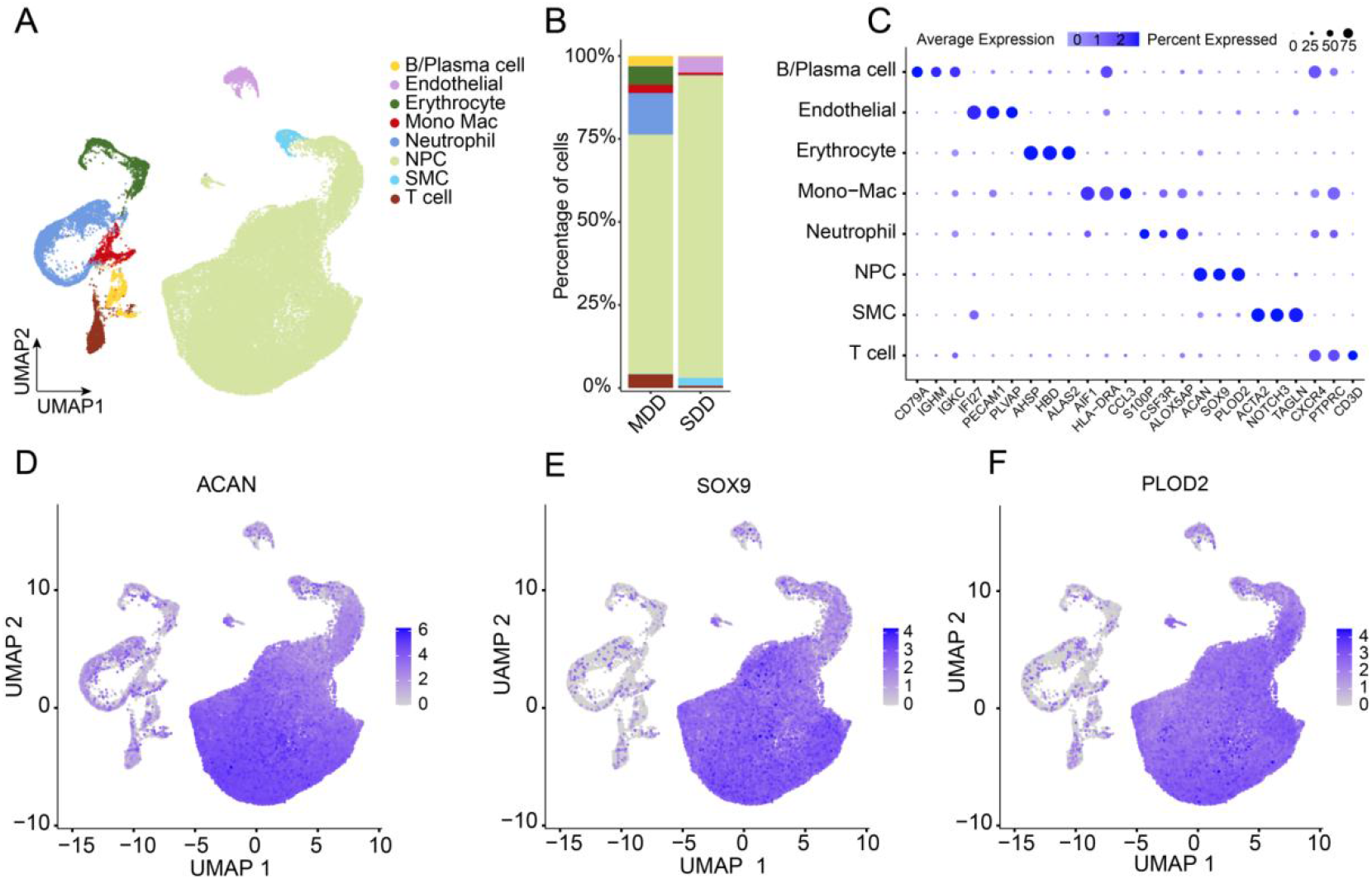
Global single-cell profiling and cell type annotation of IVDD tissues. (A) UMAP visualization of 43,637 high-quality cells derived from mild (MDD) and severe (SDD) degenerative nucleus pulposus tissues, annotated into eight major cell lineages encompassing both immune (B/plasma cells, T cells, monocytes/macrophages) and non-immune compartments (neutrophils, NPCs, endothelial cells, and erythrocytes). (B) Comparative analysis of cell type proportions between the MDD and SDD groups, highlighting an unexpected increase in the immune cell fraction within the MDD group. (C) Expression profiles of classical marker genes utilized for cell type identification, confirming robust expression of canonical markers such as (D-F) *ACAN*, *SOX9*, and *PLOD2* in the NPC population.

To demonstrate the platform’s capacity for high-resolution sub-clustering, we isolated the NPC lineage, resolving it into six distinct subpopulations (Figure 4A). Notably, this re-analysis exposed significant inter-sample heterogeneity not emphasized in the original study. For example, Met-NPCs constituted a disproportionate fraction of sample Pb-55F relative to other MDD replicates, while Fibro-NPCs were predominantly enriched in a single SDD sample, Pd-62F (Figure 4B, C). While the pan-NPC marker *ACAN* exhibited ubiquitous expression across the isolated lineage (Figure 4D), individual subsets were defined by highly specific signatures. For instance, the Fibro-NPC subpopulation was distinctly demarcated by *FBLN1*, a key regulator of collagen and extracellular matrix (ECM) organization (Figure 4H). This cluster corresponds to late-stage NPCs characterized by the upregulation of the degeneration mediator *SRGN*. Remaining NPC subtypes were accurately resolved using canonical markers, including *CHI3L2* (Figure 4E), *CP* (Figure 4F), *DKK1* (Figure 4G), and *MSMO1* (Figure 4I). Collectively, this sub-clustering workflow demonstrates how OmicsResonance bridges global cellular profiling with granular subpopulation analysis. By coupling interactive parameter tuning with transparent visual feedback, the platform empowers biologists to robustly validate literature-derived hypotheses and dynamically interrogate complex datasets to explore dataset-specific biological patterns that require independent validation.

**Figure 4.**
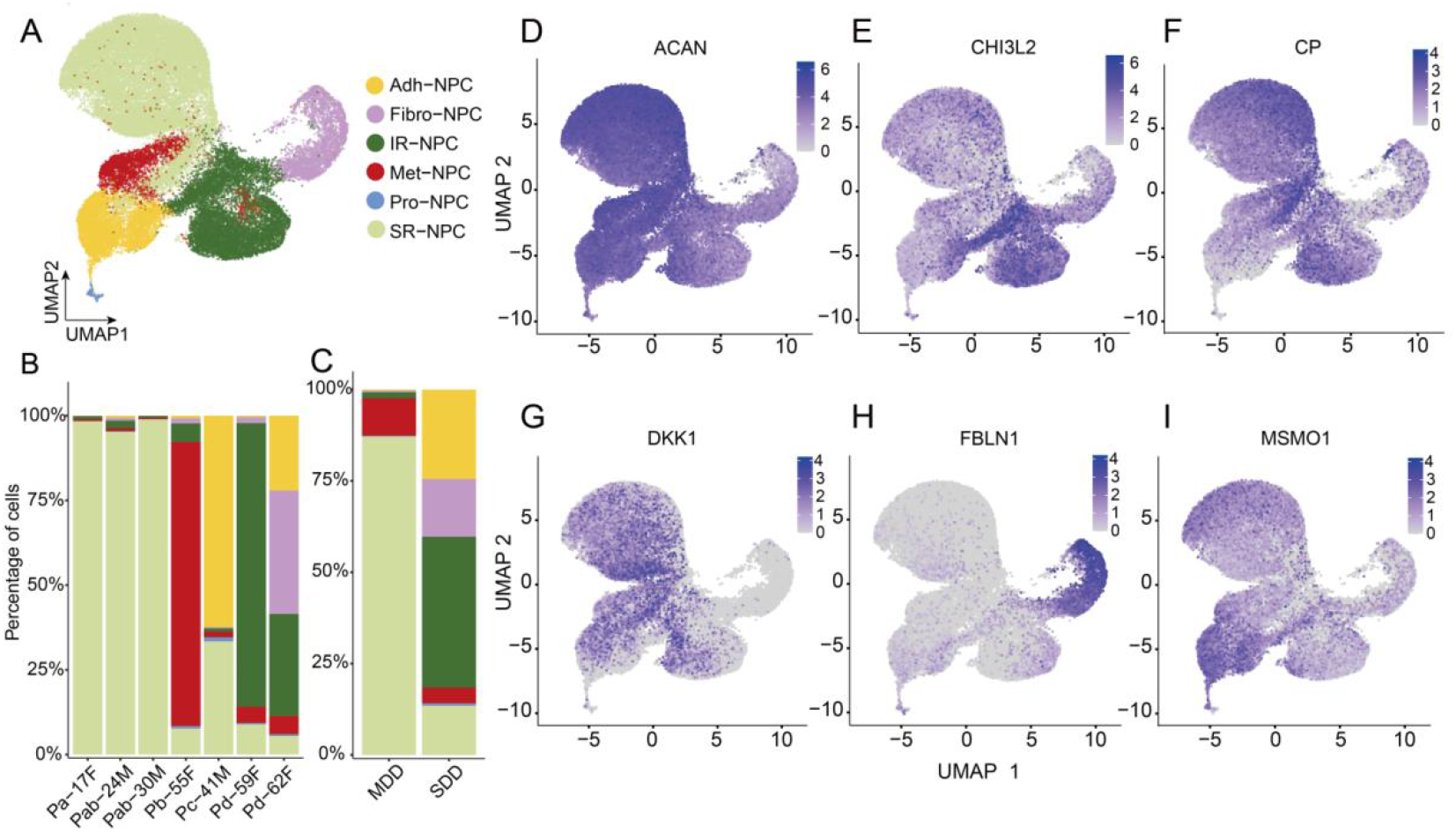
High-resolution sub-clustering and heterogeneity of nucleus pulposus cells (NPCs). (A) UMAP visualization of isolated NPCs, resolved into six distinct subpopulations. (B, C) Relative proportions of NPC subpopulations across individual biological replicates (B) and stratified by disease severity (MDD vs. SDD) (C). (D–I) UMAP feature plots illustrating the expression distributions of the pan-NPC marker *ACAN* (D) alongside specific subpopulation markers: *CHI3L2* (E), *CP* (F), *DKK1* (G), *FBLN1* (H), and *MSMO1* (I).

### Advanced Functional Profiling and Cellular State Characterization

To evaluate the platform’s capacity for multi-dimensional downstream analysis, we re-analyzed a mouse brain single-cell dataset (GSE174574)[28]. Beyond reconstructing the global cellular landscape and resolving the pro-angiogenic “healing endothelial cell” (healing EC) subpopulation (Figure 5A, B), we deployed advanced functional modules to interrogate specific cellular states and microenvironmental dynamics. To characterize the functional profile of the healing ECs, we utilized the integrated UCell[35] module, which showed elevated healing-related signature scores specifically within this subset (Figure 5C). Subsequent stemness evaluation via CytoTRACE2_1.1.0[28, 37] indicated that healing ECs exhibit the highest degree of developmental plasticity among EC subtypes (Figure 5D). Trajectory inference mapped the cellular lineage and positioned healing ECs at the initial root node of the differentiation hierarchy (Figure 5E). Finally, intercellular communication modeling via CellChat[38] predicted complex microenvironmental crosstalk, suggesting the role of healing ECs as primary signal senders in pro-inflammatory and pro-angiogenic networks (Figure 5F). Collectively, these analyses demonstrate the integrated application of multiple downstream modules within OmicsResonance. The resulting computational observations support further biological investigation but should not be interpreted as independently validated biological mechanisms.

**Figure 5.**
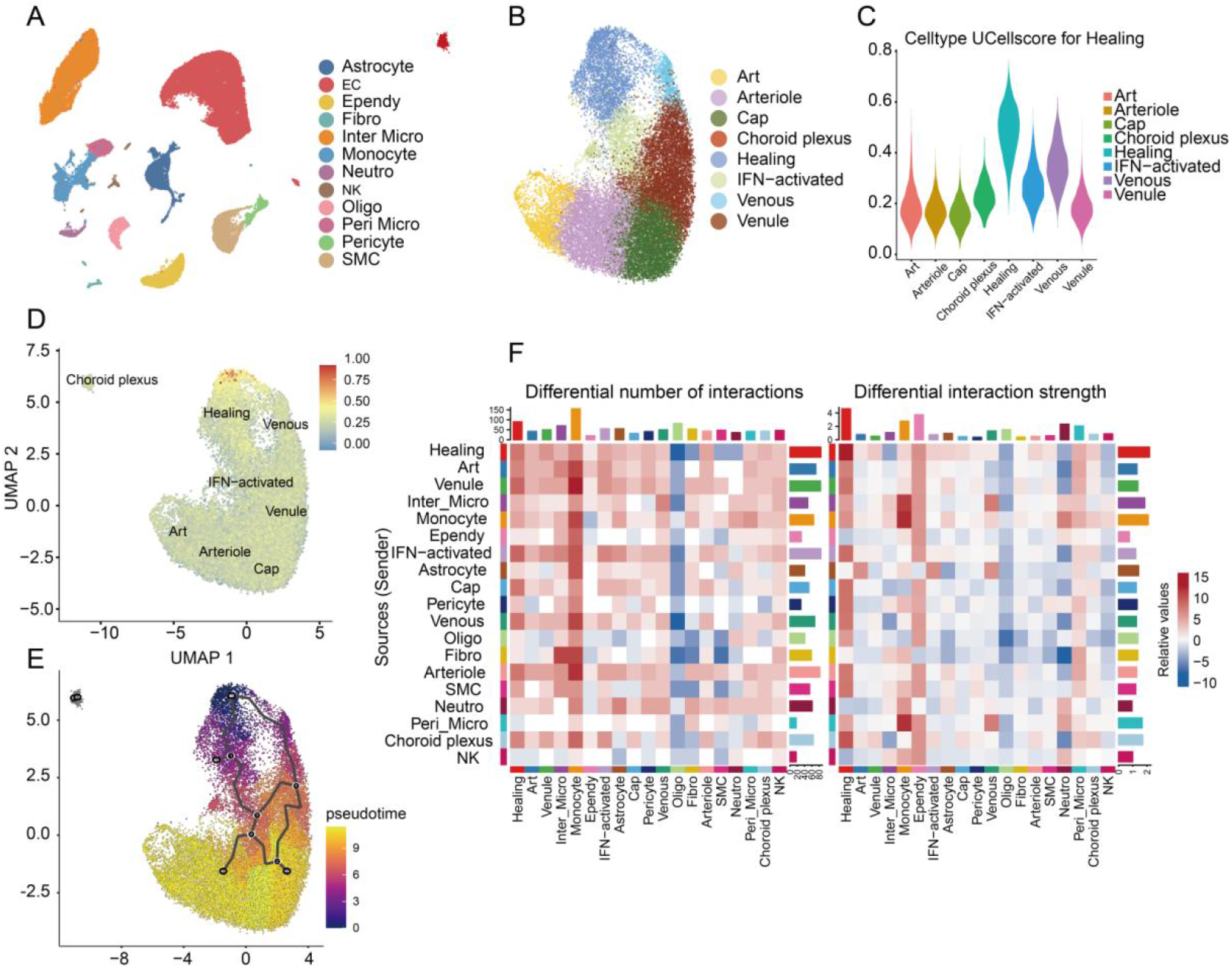
Advanced functional profiling and characterization of healing endothelial cells (ECs). (A) Global UMAP visualization of major cell populations in the mouse brain dataset (GSE174574). (B) High-resolution UMAP sub-clustering of the EC compartment, resolving the pro-angiogenic healing EC subpopulation. (C) UCell signature evaluation showing elevated healing-related scores specifically within the healing EC cluster. (D) CytoTRACE2_1.1.0 stemness assessment indicating high developmental plasticity of healing ECs relative to other subtypes. (E) Trajectory inference mapping the cellular developmental lineage, positioning healing ECs at the initial pseudotime root node. (F) CellChat analysis detailing the differential number (left) and strength (right) of intercellular communication networks among all annotated subpopulations.

## Conclusion

In conclusion, OmicsResonance offers a highly accessible and comprehensive cloud platform for single-cell transcriptomic analysis. It integrates complex bioinformatics algorithms into an intuitive web interface. This design significantly lowers the computational barriers for bench researchers. As demonstrated by our validation studies, the platform accurately replicates standard analytical baselines. It also provides reliable tools for deep functional characterization and subpopulation exploration. Importantly, OmicsResonance prioritizes analytical transparency. Features such as automated parameter archiving, interactive biological validation, and an extensible modular architecture collectively support workflow traceability and reproducibility. Ultimately, the platform assists researchers in translating high-dimensional sequencing data into robust biological insights, ensuring that the workflow is justified.

## Discussion

In this study, we introduced OmicsResonance, a comprehensive cloud ecosystem designed to democratize single-cell RNA sequencing analysis. Through systematic case studies, we demonstrated that the platform robustly handles strict quality control, multi-method cell annotation, and complex downstream functional profiling. Compared to existing web-based tools, OmicsResonance distinguishes itself by addressing critical analytical gaps. It lowers the initial technical barrier by supporting diverse input formats and one-click bilingual reporting, while simultaneously enforcing rigorous preprocessing standards, such as ambient RNA and doublet removal. Furthermore, the platform integrates a multi-method, LLM-assisted annotation workflow. In addition, we performed a quantitative comparison of the LLM-assisted annotation results with conventional annotation methods using reference cell-type annotations. The LLM-assisted workflow achieved macro-averaged F1 scores of 0.971, 0.974, and 0.929 for PBMC 3K, GSE174574, and GSE244889, respectively. It achieved the highest F1 score among the evaluated workflows for GSE174574 and GSE244889 and ranked second for PBMC 3K. However, performance was dataset dependent, and SingleR showed higher accuracy for GSE244889. These results indicate competitive practical performance without demonstrating universal superiority or the intrinsic reproducibility of independently generated LLM responses. The platform also integrates advanced analytical modules, including virtual knockout simulations and .cloupe file export, that are rarely unified in a single web service. Crucially, by automatically archiving user parameters and comprehensive execution logs, OmicsResonance supports workflow traceability and reproducibility within the centrally managed environment.

The case studies are intended to demonstrate the analytical functionality of OmicsResonance rather than to establish independently validated biological mechanisms; candidate findings require confirmation using independent datasets and, where appropriate, experimental approaches.

Several limitations of the current implementation should be acknowledged. First, OmicsResonance accepts processed expression matrices rather than raw sequencing reads. Second, computational time and memory requirements increase substantially with dataset size. Although 500,000 cells were successfully processed in the tested environment, this result should not be interpreted as a universal maximum supported dataset size. Third, cell-type annotation depends on publicly available reference databases and LLM-assisted inference and is currently optimized primarily for human and mouse datasets. Model variability, database bias, and potential hallucination cannot be completely eliminated by multi-method comparison and manual validation. The present annotation benchmark evaluated one LLM-assisted output per dataset and did not include repeated independent generations, case-by-case error analysis, or an inter-observer agreement study. In addition, a formal prospective usability study was not performed; current usability evidence is based on routine platform use, user feedback, tutorials, and example workflows rather than standardized task-completion or user-satisfaction comparisons. OmicsResonance is currently deployed on dedicated HPC infrastructure. Elastic deployment on AWS, Google Cloud, or Azure has been considered but has not yet been formally implemented or evaluated.

OmicsResonance is currently operated and maintained by Shanghai NewCore Biotechnology Co., Ltd. and is available to users without charge. Ongoing server operation, data storage, software maintenance, and technical support are supported through the company’s operating profits. Our current priority is to maintain broad and free access while monitoring user demand, computational workload, resource consumption, and operating costs. The long-term service model may be adjusted if necessary to support sustainable operation.

Future developments will extend support to spatial transcriptomics, scATAC-seq, and broader multi-omics integration. We also plan to incorporate knowledge graph integration and agent-based AI reasoning to enable automated hypothesis generation and deeper biological interpretation.

## Acknowledgement

We thank Dr. Hongsheng Li, Dr.Yuchen Cai for their extensive and critical feedback that profoundly shaped this platform, as well as the many other users for their valuable contributions. We are grateful to the members of the team for constructive discussions and suggestions. We thank the open-source community for developing and maintaining the Seurat ecosystem that form the foundation of our analysis pipeline.

## Contribution

## Author contributions

Chenhui Li: Writing – original draft, Writing – review & editing. Guicheng Zhang: Software, Validation. Lin Zeng: Conceptualization, Writing – review & editing. Mingjie Chen: Software, Writing – review & editing. Yi Lu: Software, Validation. Zhizhen Qin: Software, Data curation. Yan-kai Chen: Validation, Project administration.

## Compliance with ethics guidelines

## Ethical issues

No new human participants or animals were enrolled in this study. All analyses were based on publicly available datasets.

## Data availability and compliance statement

The datasets analyzed in this study are publicly available. The single-cell RNA sequencing datasets GSE174574 and GSE244889 were obtained from the National Center for Biotechnology Information (NCBI) Gene Expression Omnibus (GEO). The PBMC3K dataset used in the Seurat guided clustering tutorial was originally made publicly available by 10x Genomics and was accessed through the Seurat tutorial.

