## Supplementary figures and images for "OmicsResonance: An LLM-assisted cloud ecosystem for interactive and reproducible single-cell transcriptomic analysis"

### Supplemental figure S1

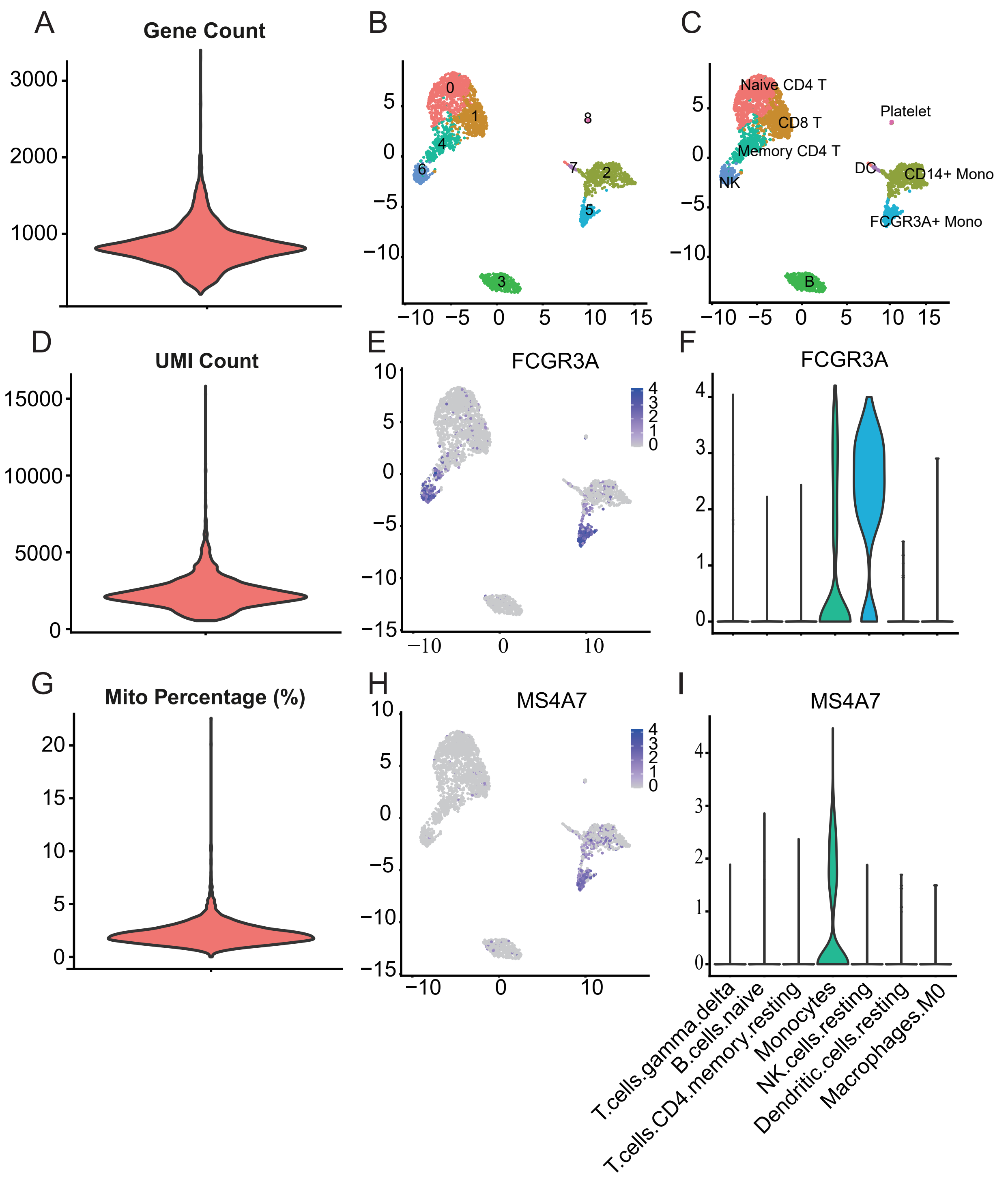
