## Supplemental figure S2 for "OmicsResonance: An LLM-assisted cloud ecosystem for interactive and reproducible single-cell transcriptomic analysis"

Supplementary Figure S2

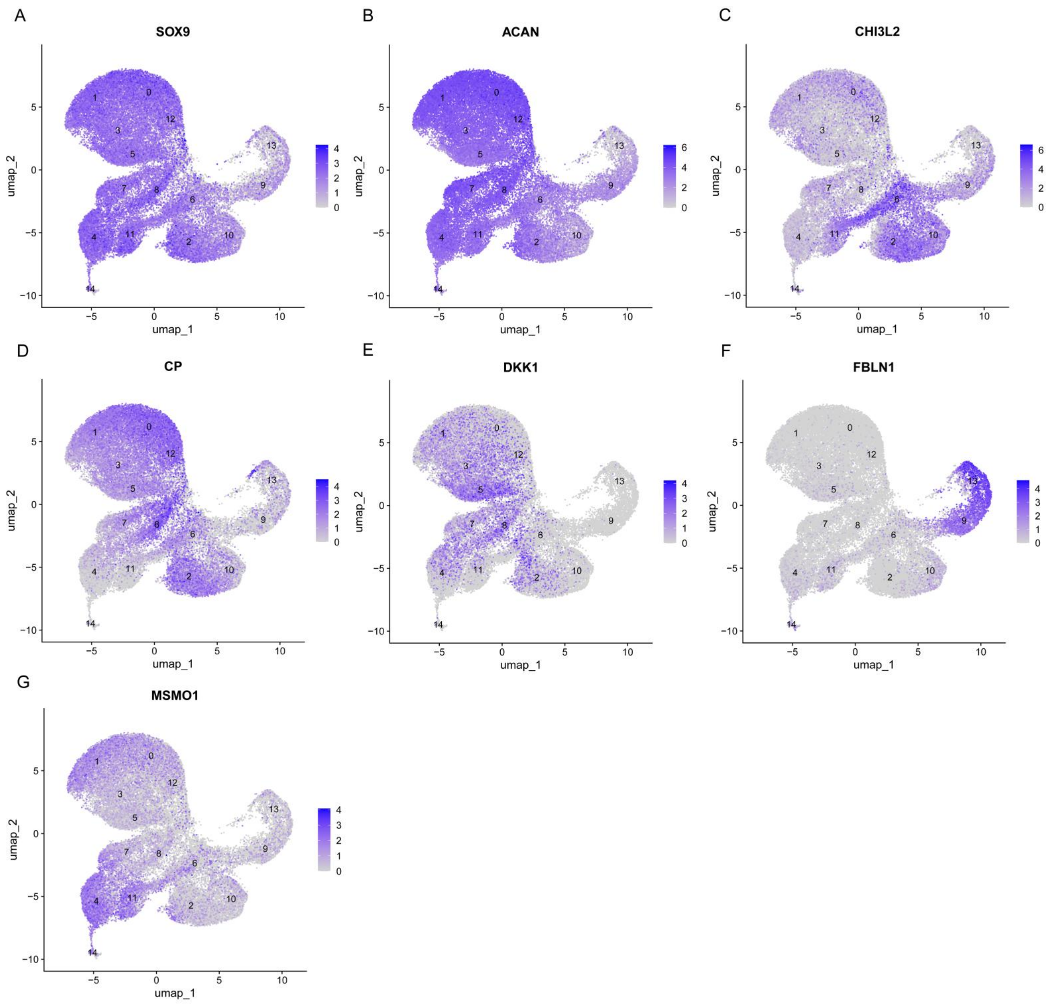


**Supplementary Figure S2: UMAP feature plots showing the distribution of gene expression in the IVDD dataset. (A) *SOX9*. (B) *ACAN*. (C) *CHI3L2*. (D) *CP*. (E) *DKK1*. (F) *FBLN1*. (G) *MSMO1*.**
